# Rethinking the efficacy of acellular pertussis vaccines for primary immunization

**DOI:** 10.1101/376947

**Authors:** M. Domenech de Cellès, P. Rohani, A. A. King

**Affiliations:** Biostatistics, Biomathematics, Pharmacoepidemiology, and Infectious Diseases (B2PHI) Unit, Institut Pasteur, Inserm U1181, University of Versailles St-Quentin-en-Yvelines, FRANCE; Odum School of Ecology, University of Georgia, Athens, GA 30602, USA; Department of Infectious Diseases, University of Georgia, Athens, GA 30602, USA; Center for the Ecology of Infectious Diseases, University of Georgia, Athens, GA 30602, USA; Department of Ecology and Evolutionary Biology, University of Michigan, Ann Arbor, MI 48109, USA; Department of Mathematics, University of Michigan, Ann Arbor, MI 48109, USA; Center for the Study of Complex Systems, University of Michigan, Ann Arbor, MI 48109, USA

## Abstract

**Background:** The US has experienced a nationwide resurgence of pertussis since the mid-1970s, despite high vaccine coverage. Short-lived immunity induced by Diphtheria-Tetanus-acellular Pertussis (DTaP) vaccines in young children is widely believed to be responsible for this growing burden. However, the duration of protection conferred by DTaP vaccines remains incompletely quantified.

**Methods and Findings:** We employed a rigorously validated, age-structured model of pertussis transmission to explore a range of hypotheses regarding the degree of waning DTaP-derived immunity. For every hypothesis, we calculated the vaccine effectiveness and the relative increase in the odds of acquiring pertussis (or odds ratio) in children aged 5 to 9 years. We then assessed the simulated DTaP vaccine traits that best reproduced the empirical values of odds ratios from recent US epidemiological studies. We found a marked association between the degree of waning immunity, the vaccine effectiveness, and the odds ratio. Unexpectedly, the odds ratio was positively associated with the vaccine effectiveness, as a consequence of non-linear, age-assortative dynamics. Based on the empirical odds ratios, we estimated that vaccine effectiveness exceeded 75% and that more than 65% of children remained immune to pertussis 5 years after the last DTaP dose.

**Conclusions:** Our results show that temporal trends in the odds of acquiring pertussis are a seriously flawed measure of the durability of vaccine-induced protection. They further demonstrate that DTaP vaccines confer imperfect, but long-lived protection. We argue that control strategies should be based upon the best available estimates of vaccine properties and the age-structure of the transmission network.

---

Pertussis, or whooping cough, is an acute respiratory disease, caused by a bacterial infection and typically characterized by a prolonged cough [1]. Despite the availability of prophylactic vaccines since the 1930s [2], recent epidemiological data indicate that the control of pertussis remains incomplete and problematic. Indeed, the disease continues to exact a heavy toll worldwide, with an estimated 24.1 [7–40] million cases and 161 [38-671] thousand deaths in 2014 in children younger than 5 yr, for the most part in low-income countries [3]. Despite large reductions in reported cases after the start of routine vaccination with Diphtheria–Tetanus–whole-cell Pertussis (DTwP, also known as DTP) vaccines, pertussis has re-emerged in several high-income countries that maintained high vaccination coverage [4, 5]. Prominently, the US has experienced a nationwide resurgence of pertussis since the mid-1970s [6, 7], with incidence highest in infants but increasing disproportionately in adolescents and adults [8, 9]. Most recent US estimates indicate that 15,737 individuals contracted pertussis in 2016, including 1,793 cases and 6 deaths in infants [10]. Additional control measures were implemented in response to this growing burden, which have met with mixed success (e.g., [11], but also [12, 13]). These difficulties illustrate both the complexity of, and knowledge gaps in, our understanding of pertussis biology and epidemiology [14, 15]. Foremost among the latter are uncertainties surrounding the nature of vaccinal immunity, which make the evaluation of vaccine efficacy in the field challenging [16].

Waning immunity following vaccination with Diphtheria-Tetanus-acellular Pertussis (DTaP) vaccines is widely believed to be responsible for the growing burden of pertussis in the US [17, 18, 19]. These subunit vaccines, based on a subset of purified antigens of *Bordetella pertussis*, were developed in response to concerns over the safety of DTwP vaccines [1]. Vaccine trials demonstrated the safety and the efficacy of DTaP vaccines [20], which progressively replaced DTwP in most high-income countries, including the US which switch to the acellular vaccines in the mid-1990s [21, 22]. However, concerns over the population-level impacts of these vaccines soon surfaced [18]. In a recent meta-analysis that included 2 case-control studies [23, 24] and 1 cohort study [25] in the US, McGirr et al. [26] estimated that the odds of acquiring pertussis increased 1.33[1.23–1.43]-fold each year since receipt of the last dose of DTaP. Similar results were obtained in another, more recent, case-control study [27]. These results have been interpreted as evidence for widespread and rapid waning of protection conferred by DTaP vaccines, casting doubt on the vaccine’s ability to control pertussis and sparking debate on the need for other control strategies [28, 17, 18] and new vaccines [29, 30]. However, it has not been established that this is a valid interpretation. Here, we demonstrate that this interpretation is in fact invalid by showing that the observed odds ratio is more consistent with much more durable vaccinal protection.

To this end, we used an empirically validated population-based model of pertussis transmission, structured according to age and parametrized using high-quality age-specific incidence data from Massachusetts [31].

As previously reported, this model successfully captured key features of pertussis epidemiology in the US (Fig. 1), chiefly the resurgence from the mid-1970s (Fig. 1B) and the concomitant shift of cases to adolescents and adults (Fig. 1C) [31]. According to this model, these changes are an end-of-honeymoon effect [32]—that is, they are the slowly manifesting but predictable consequences of incomplete historical coverage with imperfect, but nevertheless efficacious, vaccines that confer slowly waning protection and generate strong herd immunity. The underlying mechanism of this effect is illustrated in the immunological profile presented in Fig. 1A, and further explained in the legend of Fig. 1.

**Figure 1:**
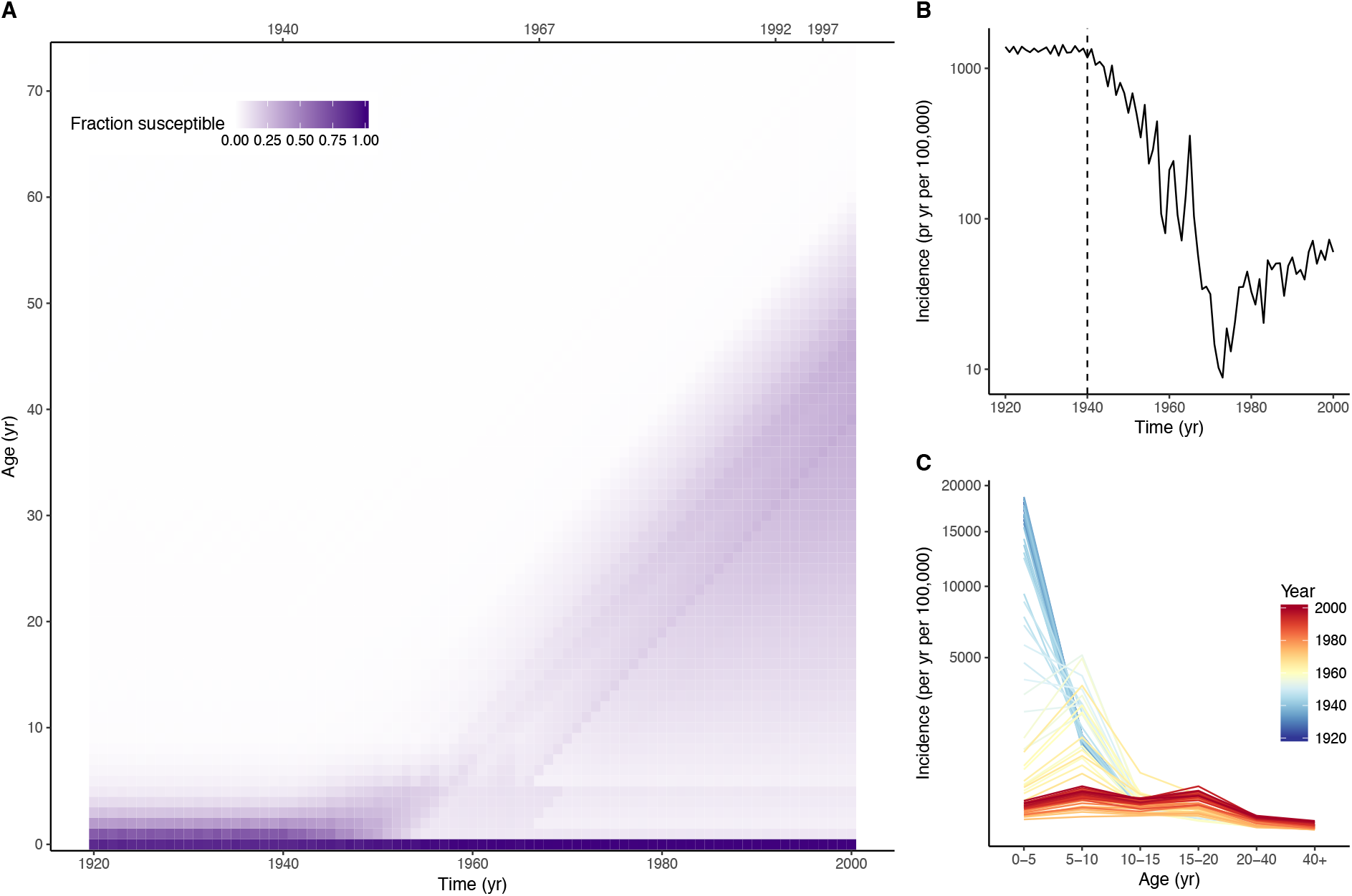
Resurgence of pertussis in the US as an end-of-honeymoon effect. This figure shows a typical simulation of a stochastic model of pertussis transmission [31], under a US-like scenario of immunization, assuming that 95% of infants are immunized with vaccines that wane slowly on average (average waning rate, 0.011 per yr, 5% probability that immunity wanes within 5 yr). A: Variations of the fraction susceptible to pertussis infection over time (*x*-axis), and according to age (*y*-axis). The top *x*-axis indicates changes in immunization practices assumed in the model: 1940, start of mass vaccination with DTwP; 1967, start of booster doses in children aged 15–18 mo (4th dose) and 4–6 yr (5th dose); 1992, start of DTaP for booster doses; 1997, start of DTaP for all doses. B: Total incidence of pertussis. The vertical dashed line indicates the assumed start time of mass vaccination with DTwP. C: Incidence of cases over age (*x*-axis) and over time (color). Each line represents a distinct year. The 3 panels illustrate the end-of-honeymoon effect, as follows. In the prevaccine era, cases are concentrated in young children who, upon recovery, develop long-lived immunity against reinfection, resulting in strong herd immunity in older individuals. The inception of mass vaccination leads to an overall reduction in transmission in those vaccinated and in the population at large. Hence, children who were not vaccinated (or in whom vaccinal protection did not initially take) are increasingly likely to reach adulthood having avoided natural infection. Concomitantly, older cohorts, with their long-lived immunity derived from natural infection during the prevaccine era, gradually die out. The result is the gradual buildup of susceptibles visible in panel A, which leads to a gradual resurgence. See Text S1 for complete details on the model formulation, parametrization, and implementation.

Unexpectedly, we also found that our model-based estimates of the increase in the odds of pertussis were consistent with those that have been obtained in the aforementioned case-control and cohort studies commonly interpreted as evidence for rapidly waning DTaP immunity (Fig. 3F in Ref. [31] and Refs. [23, 24, 25, 26, 27]). To fully resolve the apparent paradox, we took advantage of the validated model, adapting it to incorporate known changes in immunization practices in the US (including the switch to DTaP [21, 22] and the introduction of booster vaccination with Tdap in adolescents [33]) and using it to systematically compare different measures of DTaP efficacy. Specifically, we performed extensive simulations to contrast 3 measures of DTaP efficacy in 5 simulated cohorts of children born between 2001 and 2005. First, we varied the degree of waning immunity following DTaP vaccination, here quantified as the probability that DTaP-induced immunity wanes within 5 yr (*p*_5_). Second, we estimated the resulting vaccine effectiveness (VE) using standard methods. Finally, we computed the average relative yearly change in the odds of acquiring pertussis (or odds ratio, OR) after last DTaP vaccination. By comparing our model-based OR estimates with those of empirical studies [23, 24, 25, 26, 27], we sought to determine the vaccinal traits of DTaP that best explained recent epidemiological data in the US.

The results, presented in Fig. 2A, revealed a marked association between the 3 measures of vaccine efficacy. As expected, the estimated vaccine effectiveness increased as the degree of waning decreased, exceeding 90% when immunity waned in less than 15% within 5 yr. Counterintuitively, an equally strong, but *positive*, association was found between the vaccine effectiveness and the yearly increase in the odds ratio. To understand this result, we show in Fig. 2B the simulated incidence rates in children aged 5 to 9 yr (i.e., 0 to 4 yr following the last DTaP vaccination), for a range of assumptions regarding DTaP efficacy. Assuming a slowly waning, highly efficacious DTaP (*p*_5_ = 0.05, VE = 0.96), pertussis incidence was predicted to increase almost linearly over age, on average by 43% after every year since last DTaP vaccination (Fig. 2B, top panel). This result is best interpreted as a consequence of the high transmissibility of pertussis (estimated Basic Reproduction Ratio, *R*_0_ ≈ 10 in MA [31], see also Refs. [34, 35]): at vaccine coverage below the critical threshold, circulation persists and the risk of disease remains relatively high in groups with high contact rates, such as schoolchildren (Fig. S2). In contrast, the incidence profile differed markedly in the high-waning, low-efficacy DTaP scenario (Fig. 2B, lower panel). Here the incidence was predicted to peak 1-3 yr after last receipt of DTaP, resulting on average in a *decrease* in the risk of pertussis (i.e., OR ≤ 1). Under this scenario, transmissibility is so high that the pool of susceptible children—including those for whom vaccinal immunity has waned—is rapidly depleted, limiting further transmission [36]. Hence, these results demonstrate an intricate relationship between the degree of waning and the odds ratios, making their interpretation difficult and their validity as a measure of vaccine efficacy and the durability of immunity questionable.

**Figure 2:**
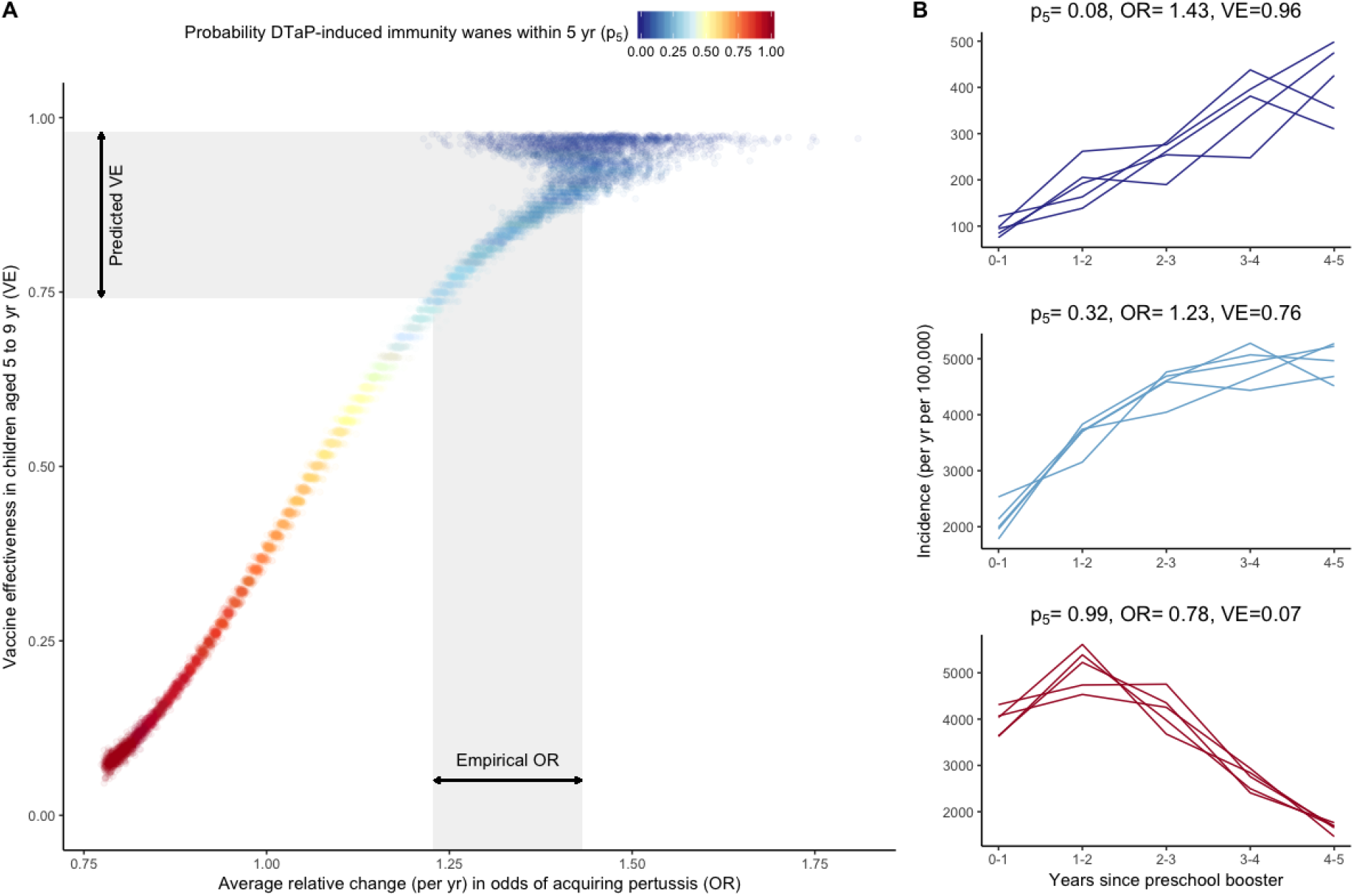
Comparison of DTaP efficacy measures. A: Association between odds ratios (x-axis) and vaccine effectiveness (y-axis), based on 10^4^ simulations with varying degrees of waning (color scale). The empirical range of odds ratios is based on the meta-analysis estimate in Ref. [26]. B: Pertussis incidence as a function of time since last receipt of DTaP for 3 values of DTaP-induced immunity waning. In each panel, the 5 lines represent 5 cohorts of children born between 2001 and 2005, tracked 0 to 4 yr after receipt of the last dose of DTaP (that is, during ages 5 to 9 yr). The *y*-axis values differ between panels, for visual clarity. See Text S1 for information on how the quantities in this figure were estimated.

Based on the meta-analysis estimate in Ref. [26] (OR=1.33 [1.23–1.43]), we predict that the effectiveness of DTaP in children aged 5 to 9 yr exceeds ≈75% (Fig. 2A). We also predict that more than 65% of children remain immune to pertussis 5 yr after the last dose of DTaP, or, equivalently, that the duration of protection exceeds approximately 12 yr. Of note, we find that the odds ratio estimates become more variable when the vaccine effectiveness exceeds 90%, as post-vaccine cases become increasingly rare and their dynamics increasingly stochastic. We propose that this finding might qualitatively explain the large estimation uncertainty found in some empirical studies [24, 25, 26], although we acknowledge other potential sources of uncertainty not incorporated into our model. To further quantify the predicted efficacy of DTaP, we also calculated the *vaccine impact*, a population-level measure of the overall reduction in transmission caused by vaccination [37, 38]. We found comparable results based on this measure, with empirical estimates of odds ratios more consistent with a vaccine impact exceeding about 50% (Fig. S4). We also found these results to be robust to alternative assumptions regarding the level of vaccine coverage, the simulation protocol, and the inclusion of demographic trends (Figs. S7–S9). Critically, these results were also insensitive to the assumed efficacy of Tdap in teenagers (Fig. S6). Altogether, we conclude from these experiments that, in stark opposition to recent claims [17, 18], recent epidemiological data in the US are actually more consistent with efficacious DTaP vaccines that confer long-term protection, reduce overall transmission, and induce herd immunity. It would be a mistake to conclude, however, that routine vaccination with DTaP alone will be sufficient to eradicate the disease [31].

With serological correlates of vaccinal protection still obscure [29], the efficacy of pertussis vaccines has been regularly debated [16]. A major point of contention remains the ability of pertussis vaccines to prevent transmission, in addition to disease [39, 40, 41, 42]. Regarding DTwP vaccines, a large body of evidence (reviewed in Ref. [15]) has shown that they can successfully reduce transmission. In contrast, there is a growing consensus that DTaP vaccines do not reduce transmission, and therefore might be inadequate to control pertussis [17, 18]. This view is partly based on evidence that DTaP generates an immune response different from that of DTwP or natural infection [43], though the immunological mechanisms of vaccinal protection remain incompletely understood. Furthermore, experimental studies in animal models have suggested that vaccination with DTaP prevents symptomatic disease, but not transmissible infection [44, 41]. We have previously argued that such results cannot be straightforwardly extrapolated to human populations inasmuch as they are inconsistent with the clear-cut signatures of herd immunity following DTaP vaccination observed in several countries [42, 15] The present findings entirely confirm this view, as they point to efficacious DTaP vaccines that confer an admittedly imperfect, but slowly waning immunity.

Our results have policy implications. First, the rationale behind future control strategies should incorporate the fact that, despite widespread belief, DTaP vaccines are actually efficacious and able to cause indirect effects via herd immunity. Second, we propose that control objectives should take into account the epidemiological dynamics of pertussis, in particular its high transmissibility. Indeed, our results suggest that a relatively high burden of pertussis—including periodic outbreaks in school-aged children—may be the norm, even with efficacious but imperfect vaccines. In view of the high transmissibility of pertussis, current DTaP vaccines are likely insufficient to eradicate the disease on their own, but they nevertheless remain an important part of effective control strategies. Empirically validated models of pertussis transmission, such as those presented here, will prove useful to define achievable control objectives, assess the impact of current control measures, and predict the effect of new control strategies. Finally, our results emphasize the complexity of pertussis epidemiology and the fact that seemingly intuitive measures of vaccine efficacy can be misleading in the face of this complexity.

## Acknowledgments

We thank M. Elizabeth Halloran for helpful comments on the manuscript. PR and AAK are supported by the National Institutes of Health (1R01AI101155) and by MIDAS, National Institute of General Medical Sciences U54-GM111274.

## Author contributions

Study conception: MDdC, PR, AAK; model development and implementation: MDdC; results analysis: MDdC, PR, AAK; writing: MDdC, PR, AAK.

## Data and code availability

All simulated data and computer codes will be deposited in a Dryad digital repository and are available upon request of the reviewers.

## References

[1] Edwards KM, Decker MD. Pertussis vaccines. In: Plotkin SA, Orenstein WA, Offit PA, editors. Vaccines. 6th ed. Philadelphia, Pa.: Elsevier Saunders; 2013. p. 447–492.

[2] Shapiro-Shapin CG. Pearl Kendrick, Grace Eldering, and the pertussis vaccine. Emerg Infect Dis. 2010 Aug;16(8):1273–8.

[3] Yeung KHT, Duclos P, Nelson EAS, Hutubessy RCW. An update of the global burden of pertussis in children younger than 5 years: a modelling study. Lancet Infect Dis. 2017 Sep;17(9):974–980.

[4] Sheridan SL, Ware RS, Grimwood K, Lambert SB. Number and order of whole cell pertussis vaccines in infancy and disease protection. JAMA. 2012 Aug;308(5):454–6.

[5] Choi YH, Campbell H, Amirthalingam G, van Hoek AJ, Miller E. Investigating the pertussis resurgence in England and Wales, and options for future control. BMC Med. 2016 Sep;14(1):121.

[6] Tanaka M, Vitek CR, Pascual FB, Bisgard KM, Tate JE, Murphy TV. Trends in pertussis among infants in the United States, 1980–1999. JAMA. 2003 Dec;290(22):2968–75.

[7] Rohani P, Drake JM. The decline and resurgence of pertussis in the US. Epidemics. 2011 Sep;3(3–4):183–8.

[8] Güriş D, Strebel PM, Bardenheier B, Brennan M, Tachdjian R, Finch E, et al. Changing epidemiology of pertussis in the United States: increasing reported incidence among adolescents and adults, 1990–1996. Clinical Infectious Diseases. 1999 Jun;28(6):1230–7.

[9] Farizo KM, Cochi SL, Zell ER, Brink EW, Wassilak SG, Patriarca PA. Epidemiological features of pertussis in the United States, 1980–1989. Clinical Infectious Diseases. 1992 Mar;14(3):708–19.

[10] Centers for Disease Control and Prevention. Provisional 2016 Reports of Notifiable Diseases. MMWR Morb Mortal Wkly Rep. 2017;65(52).

[11] Skoff TH, Blain AE, Watt J, Scherzinger K, McMahon M, Zansky SM, et al. Impact of the US Maternal Tetanus, Diphtheria, and Acellular Pertussis Vaccination Program on Preventing Pertussis in Infants <2 Months of Age: A Case-Control Evaluation. Clin Infect Dis. 2017 Sep;.

[12] Castagnini LA, Healy CM, Rench MA, Wootton SH, Munoz FM, Baker CJ. Impact of maternal postpartum tetanus and diphtheria toxoids and acellular pertussis immunization on infant pertussis infection. Clin Infect Dis. 2012 Jan;54(1):78–84.

[13] Blain AE, Lewis M, Banerjee E, Kudish K, Liko J, McGuire S, et al. An Assessment of the Cocooning Strategy for Preventing Infant Pertussis-United States, 2011. Clin Infect Dis. 2016 Dec;63(suppl 4):S221–S226.

[14] Jackson DW, Rohani P. Perplexities of pertussis: recent global epidemiological trends and their potential causes. Epidemiol Infect. 2014 Apr;142(4):672–84.

[15] Domenech de Cellès M, Magpantay FMG, King AA, Rohani P. The pertussis enigma: reconciling epidemiology, immunology and evolution. Proc Biol Sci. 2016 Jan;283(1822).

[16] Fine PE, Clarkson JA. Reflections on the efficacy of pertussis vaccines. Reviews of Infectious Diseases. 1987;9(5):866–83.

[17] Bolotin S, Harvill ET, Crowcroft NS. What to do about pertussis vaccines? Linking what we know about pertussis vaccine effectiveness, immunology and disease transmission to create a better vaccine. Pathog Dis. 2015 Nov;73(8):ftv057.

[18] Klein NP, Zerbo O. Use of acellular pertussis vaccines in the United States: can we do better? Expert Rev Vaccines. 2017 Dec;16(12):1175–1179.

[19] von König CHW. Acellular pertussis vaccines: where to go to? Lancet Infect Dis. 2017 Oct;.

[20] Zhang L, Prietsch SOM, Axelsson I, Halperin SA. Acellular vaccines for preventing whooping cough in children. Cochrane Database Syst Rev. 2012;3:CD001478.

[21] Pertussis vaccination: acellular pertussis vaccine for reinforcing and booster use–supplementary ACIP statement. Recommendations of the Immunization Practices Advisory Committee (ACIP). MMWR Recomm Rep. 1992 Feb;41(RR–1):1–10.

[22] Pertussis vaccination: use of acellular pertussis vaccines among infants and young children. Recommendations of the Advisory Committee on Immunization Practices (ACIP). MMWR Recomm Rep. 1997 Mar;46(RR–7):1–25.

[23] Klein NP, Bartlett J, Rowhani-Rahbar A, Fireman B, Baxter R. Waning protection after fifth dose of acellular pertussis vaccine in children. N Engl J Med. 2012 Sep;367(11):1012–9.

[24] Misegades LK, Winter K, Harriman K, Talarico J, Messonnier NE, Clark TA, et al. Association of childhood pertussis with receipt of 5 doses of pertussis vaccine by time since last vaccine dose, California, 2010. JAMA. 2012 Nov;308(20):2126–32.

[25] Tartof SY, Lewis M, Kenyon C, White K, Osborn A, Liko J, et al. Waning immunity to pertussis following 5 doses of DTaP. Pediatrics. 2013 Apr;131(4):e1047–52.

[26] McGirr A, Fisman DN. Duration of pertussis immunity after DTaP immunization: a meta-analysis. Pediatrics. 2015 Feb;135(2):331–43.

[27] Klein NP, Bartlett J, Fireman B, Aukes L, Buck PO, Krishnarajah G, et al. Waning protection following 5 doses of a 3-component diphtheria, tetanus, and acellular pertussis vaccine. Vaccine. 2017 Jun;35(26): 3395–3400.

[28] Burns DL, Meade BD, Messionnier NE. Pertussis resurgence: perspectives from the Working Group Meeting on pertussis on the causes, possible paths forward, and gaps in our knowledge. J Infect Dis. 2014 Apr;209 Suppl 1:S32–S35.

[29] Mills KHG, Ross PJ, Allen AC, Wilk MM. Do we need a new vaccine to control the re-emergence of pertussis? Trends Microbiol. 2014 Feb;22(2):49–52.

[30] Clark TA, Messonnier NE, Hadler SC. Pertussis control: time for something new? Trends Microbiol. 2012 May;20(5):211–213.

[31] Domenech de Cellès M, Magpantay FMG, King AA, Rohani P. The impact of past vaccination coverage and immunity on pertussis resurgence. Sci Transl Med. 2018 Mar;10(434).

[32] McLean AR, Anderson RM. Measles in developing countries. Part II. The predicted impact of mass vaccination. Epidemiology and Infection. 1988 Jun;100(3):419–42.

[33] Broder KR, Cortese MM, Iskander JK, Kretsinger K, Slade BA, Brown KH, et al. Preventing tetanus, diphtheria, and pertussis among adolescents: use of tetanus toxoid, reduced diphtheria toxoid and acellular pertussis vaccines recommendations of the Advisory Committee on Immunization Practices (ACIP). MMWR Recomm Rep. 2006 Mar;55(RR–3):1–34.

[34] Gambhir M, Clark TA, Cauchemez S, Tartof SY, Swerdlow DL, Ferguson NM. A change in vaccine efficacy and duration of protection explains recent rises in pertussis incidence in the United States. PLoS Comput Biol. 2015 Apr;11(4):e1004138.

[35] Rohani P, Zhong X, King AA. Contact network structure explains the changing epidemiology of pertussis. Science. 2010 Nov;330(6006):982–5.

[36] Keeling MJ, Rohani P. Modeling infectious diseases in humans and animals. Princeton: Princeton University Press; 2008. Available from: http://www.loc.gov/catdir/toc/fy0805/2006939548.html.

[37] Halloran ME, Longini IM, Struchiner CJ. Design and analysis of vaccine studies. Statistics for biology and health. New York: Springer; 2010.

[38] Magpantay FMG, Riolo MA, Domenech de Cellès M, King AA, Rohani P. Epidemiological consequences of imperfect vaccines for immunizing infections. SIAM J Appl Math. 2014;74(6):1810–1830.

[39] Fine PE, Clarkson JA. The recurrence of whooping cough: possible implications for assessment of vaccine efficacy. Lancet. 1982 Mar;1(8273):666–9.

[40] Rohani P, Earn DJ, Grenfell BT. Impact of immunisation on pertussis transmission in England and Wales. Lancet. 2000 Jan;355(9200):285–6.

[41] Warfel JM, Zimmerman LI, Merkel TJ. Acellular pertussis vaccines protect against disease but fail to prevent infection and transmission in a nonhuman primate model. Proc Natl Acad Sci USA. 2014 Jan;111(2):787–92.

[42] Domenech de Cellès M, Riolo MA, Magpantay FMG, Rohani P, King AA. Epidemiological evidence for herd immunity induced by acellular pertussis vaccines. Proc Natl Acad Sci USA. 2014 Feb;111(7):E716–7.

[43] Ryan M, Murphy G, Ryan E, Nilsson L, Shackley F, Gothefors L, et al. Distinct T-cell subtypes induced with whole cell and acellular pertussis vaccines in children. Immunology. 1998 Jan;93(1):1–10.

[44] Smallridge WE, Rolin OY, Jacobs NT, Harvill ET. Different Effects of Whole-Cell and Acellular Vaccines on *Bordetella* Transmission. J Infect Dis. 2014 Mar;.

